# OnACID: Online Analysis of Calcium Imaging Data in Real Time

**DOI:** 10.1101/193383

**Authors:** Andrea Giovannucci, Johannes Friedrich, Matt Kaufman, Anne Churchland, Dmitri Chklovskii, Liam Paninski, Eftychios A. Pnevmatikakis

**Affiliations:** Center for Computational Biology, Flatiron Institute, Simons Foundation, New York, NY 10010; Department of Statistics, Center for Theoretical Neuroscience, Columbia University, New York, NY 10027; Cold Spring Harbor Laboratory, Cold Spring Harbor, NY 11724

**Author notes:** Rough draft of work to appear at NIPS 2017. These authors contributed equally to this work (Andrea Giovannucci, Johannes Friedrich).

## Abstract

Optical imaging methods using calcium indicators are critical for monitoring the activity of large neuronal populations in vivo. Imaging experiments typically generate a large amount of data that needs to be processed to extract the activity of the imaged neuronal sources. While deriving such processing algorithms is an active area of research, most existing methods require the processing of large amounts of data at a time, rendering them vulnerable to the volume of the recorded data, and preventing realtime experimental interrogation. Here we introduce OnACID, an Online framework for the Analysis of streaming Calcium Imaging Data, including i) motion artifact correction, ii) neuronal source extraction, and iii) activity denoising and deconvolution. Our approach combines and extends previous work on online dictionary learning and calcium imaging data analysis, to deliver an automated pipeline that can discover and track the activity of hundreds of cells in real time, thereby enabling new types of closed-loop experiments. We apply our algorithm on two large scale experimental datasets, benchmark its performance on manually annotated data, and show that it outperforms a popular offline approach.

## 1 Introduction

Calcium imaging methods continue to gain traction among experimental neuroscientists due to their capability of monitoring large targeted neuronal populations across multiple days or weeks with decisecond temporal and single-neuron spatial resolution. To infer the neural population activity from the raw imaging data, an analysis pipeline is employed which typically involves solving the following problems (all of which are still areas of active research): i) correcting for motion artifacts during the imaging experiment, ii) identifying/extracting the sources (neurons and axonal or dendritic processes) in the imaged field of view (FOV), and iii) denoising and deconvolving the neural activity from the dynamics of the expressed calcium indicator.

The fine spatiotemporal resolution of calcium imaging comes at a data rate cost; a typical two-photon (2p) experiment on a 512×512 pixel large FOV imaged at 30Hz, generates ∼50GB of data (in 16-bit integer format) per hour. These rates can be significantly higher for other planar and volumetric imaging techniques, e.g., light-sheet [1] or SCAPE imaging [4], where the data rates can exceed 1TB per hour. The resulting data deluge poses a significant challenge.

Of the three basic pre-processing problems described above, the problem of source extraction faces the most severe scalability issues. Popular approaches reshape the data movies into a large array with dimensions (#pixels) x (#timesteps), that is then factorized (e.g., via independent component analysis [22] or constrained non-negative matrix factorization (CNMF) [28]) to produce the locations in the FOV and temporal activities of the imaged sources. While effective for small or medium datasets, direct factorization can be impractical, since a typical experiment can quickly produce datasets larger than the available RAM. Several strategies have been proposed to enhance scalability, including parallel processing [9], spatiotemporal decimation [10], dimensionality reduction [25], and out-of-core processing [14]. While these approaches enable efficient processing of larger datasets, they still require significant storage, power, time, and memory resources.

Apart from recording large neural populations, optical methods can also be used for stimulation [5]. Combining optogenetic methods for recording and perturbing neural ensembles opens the door to exciting closed-loop experiments [26, 16, 8], where the pattern of the stimulation can be determined based on the recorded activity during behavior. In a typical closed-loop experiment, the monitored/perturbed regions of interest (ROIs) have been preselected by analyzing offline a previous dataset from the same FOV. Monitoring the activity of a ROI, which usually corresponds to a soma, typically entails averaging the fluorescence over the corresponding ROI, resulting in a signal that is only a proxy for the actual neural activity and which can be sensitive to motion artifacts and drifts, as well as spatially overlapping sources, background/neuropil contamination, and noise. Furthermore, by preselecting the ROIs, the experimenter is unable to detect and incorporate new sources that become active later during the experiment, which prevents the execution of truly closed-loop experiments.

In this paper, we present an Online, single-pass, algorithmic framework for the Analysis of Calcium Imaging Data (OnACID). Our framework is highly scalable with minimal memory requirements, as it processes the data in a streaming fashion one frame at a time, while keeping in memory a set of low dimensional sufficient statistics and a small minibatch of the last data frames. Every frame is processed in four sequential steps: i) The frame is registered against the previous denoised (and registered) frame to correct for motion artifacts. ii) The fluorescence activity of the already detected sources is tracked. iii) Newly appearing neurons and processes are detected and incorporated to the set of existing sources. iv) The fluorescence trace of each source is denoised and deconvolved to provide an estimate of the underlying spiking activity.

Our algorithm integrates and extends the online NMF algorithm of [21], the CNMF source extraction algorithm of [28], and the near-online deconvolution algorithm of [11], to provide a framework capable of real time identification and processing of hundreds of neurons in a typical 2p experiment (512x512 pixel wide FOV imaged at 30Hz), enabling novel designs of closed-loop experiments.

We apply OnACID to two large-scale (50 and 65 minute long) mouse *in vivo* 2p datasets; our algorithm can find and track hundreds of neurons faster than real-time, and outperforms the CNMF algorithm of [28] benchmarked on multiple manual annotations using a precision-recall framework.

## 2 Methods

We illustrate OnACID in process in Fig. 1. At the beginning of the experiment (Fig. 1-left), only a few components are active, as shown in the panel A by the max-correlation image^1^, and these are detected by the algorithm (Fig. 1B). As the experiment proceeds more neurons activate and are subsequently detected by OnACID (Fig. 1 middle, right) which also tracks their activity across time (Fig. 1C). See also Supplementary Movie 1 for an example in simulated data.

**Figure 1:**
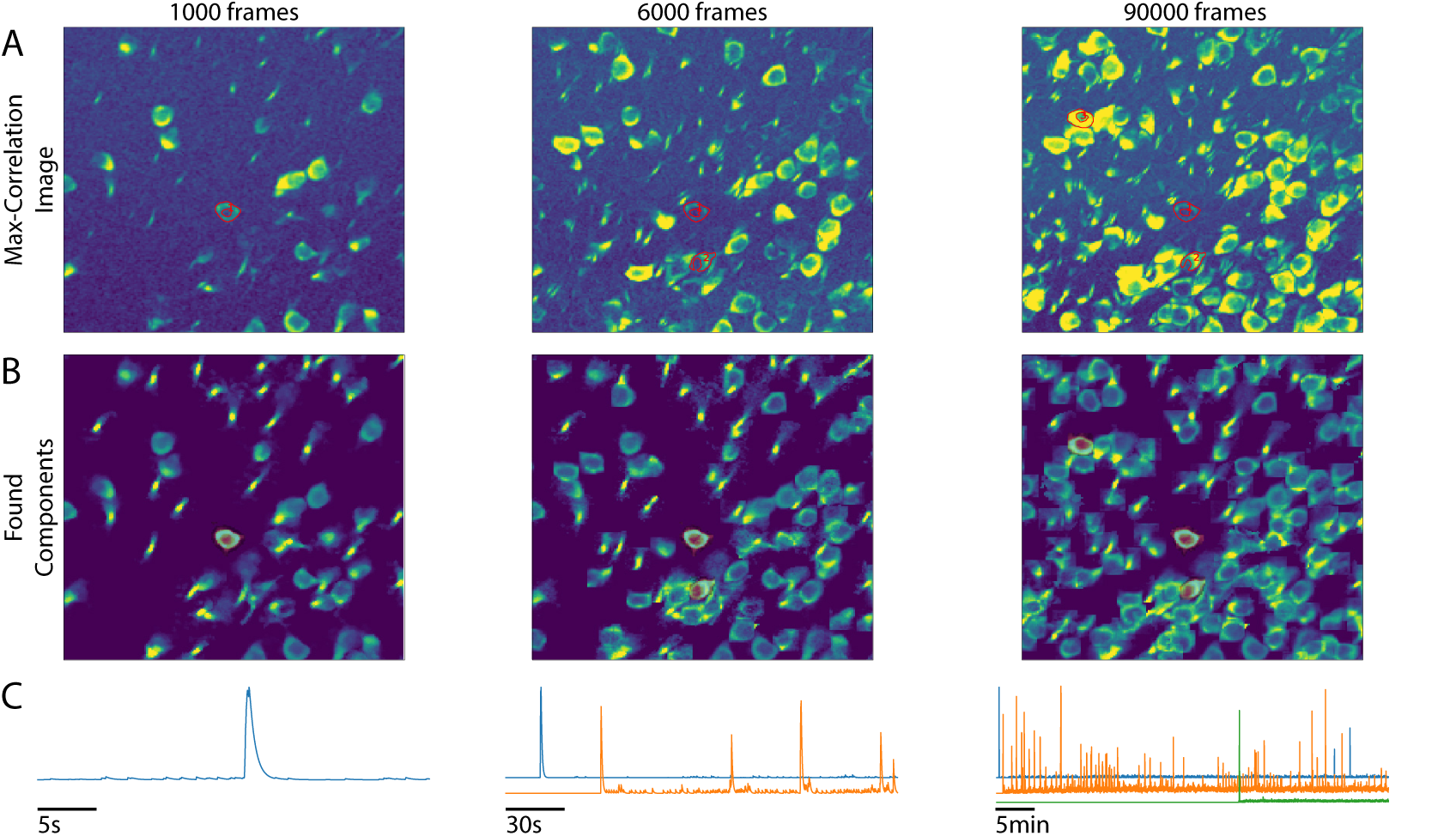
Illustration of the online data analysis process. Snapshots of the online analysis after processing 1000 frames (left), 6000 frames (middle), and 90000 frames (right). A) "Max-correlation" image of registered data at each snapshot point (see text for definition). B) Components found by OnACID up to each point. C) Examples of neuron activity traces (marked by contours in panel A and highlighted in red in panel B). As the experiment proceeds, OnACID detects newly active neurons and tracks their activity.

Next, we present the steps of OnACID in more detail.

**Motion correction:** Our online approach allows us to employ a very simple yet effective motion correction scheme: each *denoised* dataframe can be used to register the next incoming noisy dataframe. To enhance robustness we use the denoised background/neuropil signal (defined in the later section) as a template to align the next dataframe. We use rigid, sub-pixel registration [17], although piecewise rigid registration can also be used at an additional computational cost. This simple alignment process is not suitable for offline algorithms due to noise in the raw data, leading to the development of various algorithms based on template matching [15,25,27] or Hidden Markov Models [7,19].

**Source extraction:** A standard approach for source extraction is to model the fluorescence within a matrix factorization framework [22, 28]. Let *Y* ϵ ℝ^*dxT*^ denote the observed fluorescence across space and time in a matrix format, where d denotes the number of imaged pixels, and *T* the length of the experiment in timepoints. If the number of imaged sources is *K*, then let *A* ϵ ℝ^*dxK*^ denote the matrix where column i encodes the "spatial footprint" of the source *i*. Similarly, let *C* ϵ ℝ^*K*x*T*^ denote the matrix where each row encodes the temporal activity of the corresponding source. The observed data matrix can then be expressed as

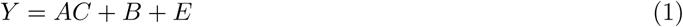

where *B*, *E* ϵ ℝ^*d*x*T*^ denote matrices for background/neuropil activity and observation noise, respectively. A common approach, introduced in [28], is to express the background matrix B as a low rank matrix, i.e., *B* = *bf*, where *b* ϵ ℝ^*d*x*nb*^ and *f* ϵ ℝ^*nb*x*T*^ denote the spatial and temporal components of the low rank background signal, and *n*_*b*_ is a small integer, e.g., *n*_*b*_ = 1, 2. The CNMF framework of [28] operates by alternating optimization of [*A*, *b*] given the data *Y*. and estimates of [*C*; *f*], and vice versa, where each column of A is constrained to be zero outside of a neighborhood around its previous estimate. This strategy exploits the spatial locality of each neuron to reduce the computational complexity. This framework can be adapted to a data streaming setup using the online NMF algorithm of [21], where at the observed fluorescence at time *t* can be written as

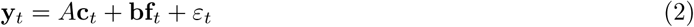

Proceeding in a similar alternating way, the activity of all neurons at time *t*, *c*_*t*_, and temporal background f_*t*_, given y_*t*_ and the spatial footprints and background [*A*, *b*], can be found by solving a nonnegative least squares problem, whereas [*A*, *b*] can be estimated efficiently as in [21] by only keeping in memory the sufficient statistics (where we define *c*_*t*_ = [c_t_; f_t_])

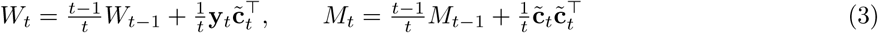

while at the same time enforcing the same spatial locality constraints as in the CNMF framework.

**Deconvolution:** The online framework presented above estimates the demixed fluorescence traces c^1^,…, c^*K*^ of each neuronal source. The fluorescence is a filtered version of the underlying neural activity that we want to infer. To further denoise and deconvolve the neural activity from the dynamics of the indicator we use the OASIS algorithm [11] that implements the popular spike deconvolution algorithm of [31] in a nearly online fashion by adapting the highly efficient Pool Adjacent Violators Algorithm used in isotonic regression[3]. The calcium dynamics is modeled with a stable autoregressive process of order *p*, 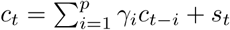 We use *p* = 1 here, but can extend to *p* =2 to incorporate the indicator rise time [11]. OASIS solves a modified LASSO problem

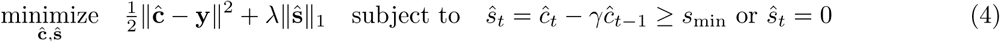

where the ℓ_1_ penalty on ŝ or the minimal spike size *s*_min_ can be used to enforce sparsity of the neural activity. The algorithm progresses through each time series sequentially from beginning to end and backtracks only to the most recent spike. We can further restrict the lag to few frames, to obtain a good approximate solution applicable for real-time experiments.

**Detecting new components:** The approach explained above enables tracking the activity of a fixed number of sources, and will ignore neurons that become active later in the experiment. To account for a variable number of sources in an online NMF setting, [12] proposes to add a new random component when the correlation coefficient between each data frame and its representation in terms of the current factors is lower than a threshold. This approach is insufficient here since the footprint of a new neuron in the whole FOV is typically too small to modify the correlation coefficient significantly.

We approach the problem by introducing a buffer that contains the last *l*_*b*_instances of the residual signal *r*_t_ = y_*t*_—A*c*_t_—bf_*t*_, where *l*_*b*_is a reasonably small number, e.g., *l*_*b*_= 100. On this buffer, similarly to [28], we perform spatial smoothing with a Gaussian kernel with radius similar to the expected neuron radius, and then search for the point in space that explains the maximum variance. New candidate components a_new_, and c_new_are estimated by performing a local rank-1 NMF of the residual matrix restricted to a fixed neighborhood around the point of maximal variance.

To limit false positives, the candidate component is screened for quality. First, to prevent noise overfitting, the shape a_new_must be significantly correlated (e.g., r ∼ 0.8 — 0.9) to the residual buffer averaged over time and restricted to the spatial extent of anew. Moreover, if anew significantly overlaps with any of the existing components, then its temporal component cnew must not be highly correlated with the corresponding temporal components; otherwise we reject it as a possible duplicate of an existing component. Once a new component is accepted, [A, b], [C; f] are augmented with a_new_and c_new_respectively, and the sufficient statistics are updated as follows:

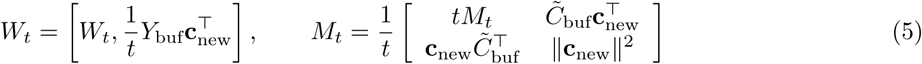

where Y_buf_, 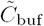 denote the matrices Y, [C; f], restricted to the last N frames that the buffer stores. This process is repeated until no new components are accepted, at which point the next frame is read and processed. The whole online procedure is described in Algorithm 1; the supplement includes pseudocode description of the referenced routines.

**Initialization:** To initialize our algorithm we use the CNMF algorithm on a short initial batch of data of length T_b_, (e.g., T_b_= 1000). The sufficient statistics are initialized from the components that the offline algorithm finds according to (3). To ensure that new components are also initialized in the darker parts of the FOV, each data frame is normalized with the (running) mean for every pixel, during both the offline and the online phases.

### Algorithm 1 OnACID

**Require:** Data matrix Y, initial estimates A, b,C, f, S, current number of components K, current timestep

*t*´ rest of parameters.

1: *W* = Y [:; 1 : t ′]C^T^′ / *t*′

2: *M* = *CC*^*T*^′ / *t*′ ⊳ Initialize sufficient statistics

3: *G* = DETERMINEGROUPS([A; b];*K*) .⊳ Alg. S1-S2

4: R_buf_ = [*Y* − [A; b][C; *f*]][:; t′ − *l*_*b*_ + 1 : t′] .⊳ Initialize residual buffer

5: *t* = *t*′

6: **while** there is more data **do**

7: *t*←*t* + 1

8: *y*_*t*_←ALIGNFRAME (y_*t*_; **bf**_*t*-1_) .⊳ [17]

9: [c_t_; f_t_] ←UPDATE TRACES([A; b]; [c_*t*-1_; f_*t*-1_]; y_*t*_; *G*) .⊳ Alg. S3

10: C; S← OASIS(C γ s_min_λ) .⊳ [11]

11: [A; b]; [C; f];K; G;R_buf_,W;M ←

12: DETECTNEWCOMPONENTS([A; b]; [C; *f*];K; G;R_buf_, y_*t*_;W;M) .⊳ Alg. S4

13: **if**mod (t − t′; T_p_) = 0 then .⊳ Update W;M; [A; b] every T_p_timesteps

14: *W*;*M* ←UPDATESUFFSTATISTICS(*W*;*M*; y_*t*_[*c*_*t*_; f_t_]) .⊳Equation (3)

15: [A; b] ←UPDATESHAPES[W;M; [A; b]]. ⊳Alg. S5

16: **end if**

17: **end while**

18: **return** A,b,*C*,f, *S*

**Algorithmic Speedups**: Several algorithmic and computational schemes are employed to boost the speed of the algorithm and make it applicable to real-time large-scale experiments. In [21] block coordinate descent is used to update the factors A, warm started at the value from the previous iteration. The same trick is used here not only for A, but also for C, since the calcium traces are continuous and typically change slowly. Moreover, the temporal traces of components that do not spatially overlap with each other can be updated simultaneously in vector form; we use a simple greedy scheme to partition the components into spatially non-overlapping groups.

Since neurons’ shapes are not expected to change at a fast timescale, updating their values (i.e., recomputing A and b) is not required at every timepoint; in practice we update every 100 timesteps. Additionally, the sufficient statistics W_t_, M_t_are only needed for updating the estimates of [A, b] so they can be updated only when required. Motion correction can be sped up by estimating the motion only on a small (active) contiguous part of the FOV. Finally, as shown in [10], spatial decimation can bring significant speed benefits without compromising the quality of the results.

### Software

OnACID is implemented in Python and is available at <u>https://github.com/simonsfoundation/caiman</u> as part of the CaImAn package [14].

## 3 Results

**Benchmarking on simulated data**: To compare to ground truth spike trains, we simulated a 2000 frame dataset taken at 30Hz over a 256 x 256 pixel wide FOV containing 400 "donut" shaped neurons with Poisson spike trains (see supplement for details). OnACID was initialized on the first 500 frames. During initialization, 265 active sources were accurately detected (Fig. S2). After the full 2000 frames, the algorithm had detected and tracked all active sources, plus one false positive (Fig. 2A).

**Figure 2:**
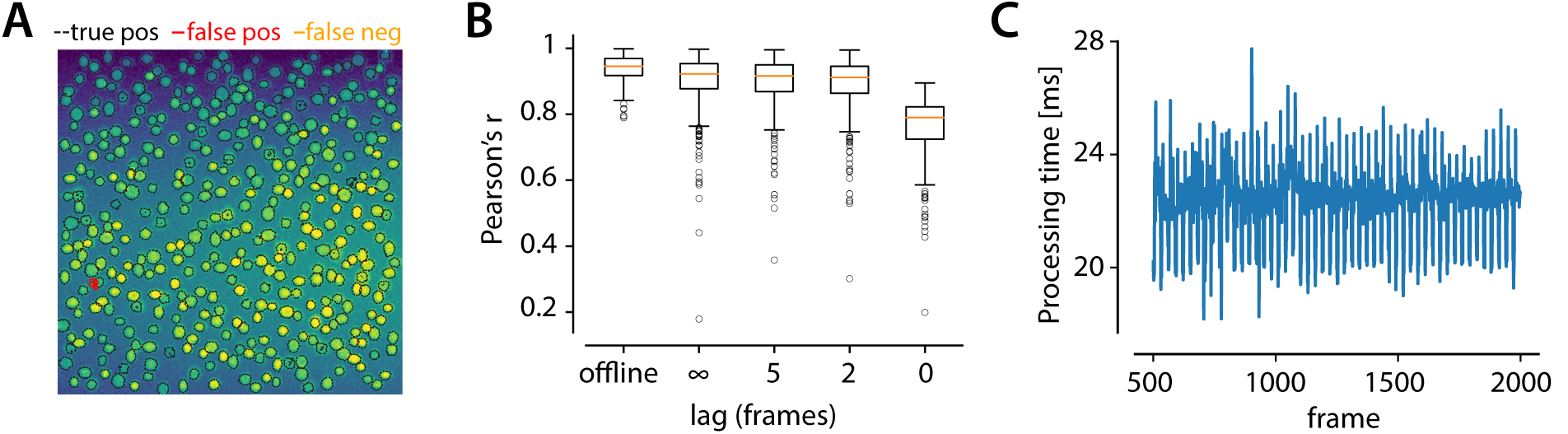
Application to simulated data. A) Detected and missed components. False positives are due to the processing of the initial batch (cf. Fig. S2). B) Tukey boxplot of spike train correlations with ground truth. Online deconvolution recovers spike trains well and the accuracy increases with the allowed lag in spike assignment. C) Processing time is less than 33 ms for all the frames.

After detecting a neuron, we need to extract its spikes with a short time-lag, to enable interesting closed loop experiments. To quantify performance we measured the correlation of the inferred spike train with the ground truth (Fig. 2B). We varied the lag in the online estimator, i.e. the number of future samples observed before assigning a spike at time zero. Lags of 2-5 yield already similar results as the solution with unrestricted lag. A further requirement for online closed-loop experiments is that the computational processing is fast enough. To balance the computational load over frames, we distributed here the shape update over the frames, while still updating each neuron every 30 frames on average. Because the shape update is the last step of the loop in Algorithm 1, we keep track of the time already spent in the iteration and increase or decrease the number of updated neurons accordingly. In this way the frame processing rate remained always higher than 30Hz (Fig. 2C).

**Application to** *in vivo* **2p mouse hippocampal data**: Next we considered a larger scale (90K frames, 480 x 480 pixels) real 2p calcium imaging dataset taken at 30Hz (i.e., 50 minute experiment). Motion artifacts were corrected prior to the analysis described below. The online algorithm was initialized on the first 1000 frames of the dataset using a Python implementation of the CNMF algorithm found in the CaImAn package [14]. During initialization 139 active sources were detected; by the end of all 90K frames, 727 active sources had been detected and tracked (5 of which were discarded due to their small size).

**Benchmarking against offline processing and manual annotations**: We collected manual annotations from two independent labelers who were instructed to find round or donut shaped neurons of similar size using the ImageJ Cell Magic Wand tool [32] given i) a movie obtained by removing a running 20th percentile (as a crude background approximation) and downsampling in time by a factor of 10, and ii) the max-correlation image. The goal of this pre-processing was to suppress silent and promote active cells. The labelers found respectively 872 and 880 ROIs. We also compared with the CNMF algorithm applied to the whole dataset which found 904 sources (805 after filtering for size).

To quantify performance we used a precision/recall framework similar to [2]. As a distance metric between two cells we used the Jaccard distance, and the pairing between different annotations was computed using the Hungarian algorithm, where matches with distance > 0.7 were discarded^2^. Table 1 summarizes the results within the precision/recall framework. The online algorithm not only matches but outperforms the offline approach of CNMF, reaching high performance values (Fi = 0.79 and 0.78 against the two manual annotations, as opposed to 0.71 against both annotations for CNMF). The two annotations matched closely with each other (F_1_= 0.89), indicating high reliability, whereas OnACID vs CNMF also produced a high score (Fi = 0.79), suggesting significant overlap in the mismatches between the two algorithms against manual annotations.

**Table 1:**
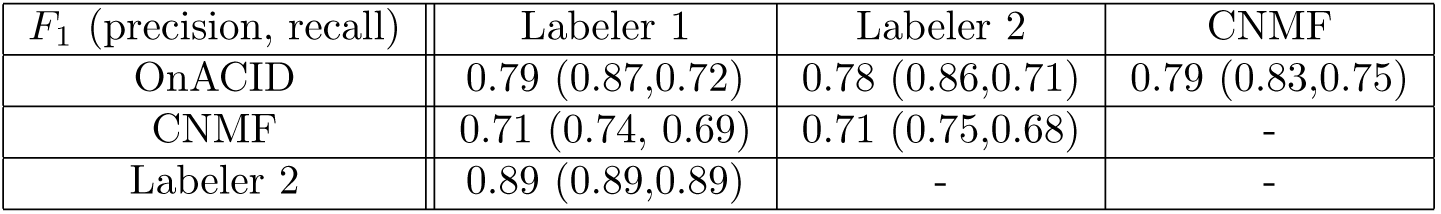
OnACID significantly outperforms the offline CNMF approach. Benchmarking is against two independent manual annotations within the precision/recall (and their harmonic mean *F*_1_ score) framework. For each row-column pair, the column dataset is regarded as the ground truth.

Fig. 3 offers a more detailed view, where contour plots of the detected components are superimposed on the max-correlation image for the online (Fig. 3A) and offline (Fig. 3B) algorithms (white) and the annotations of Labeler 1 (red) restricted to a 200x200 pixel part of the FOV. Annotations of matches and mismatches between the online algorithm and the two labelers, as well as between the two labelers in the entire FOV are shown in Figs. S3-S8. For the automated procedures binary masks and contour plots were constructed by thresholding the spatial footprint of each component at a level equal to 0.2 times its maximum value. A close inspection at the matches between the online algorithm and the manual annotation (Fig. 3A-left) indicates that neurons with a strong footprint in the max-correlation image (indicating calcium transients with high amplitude compared to noise and background/neuropil activity) are reliably detected, despite the high neuron density and level of overlap. On the other hand, mismatches (Fig. 3B-left) can sometimes be attributed to shape mismatches, manually selected components with no signature in the max-correlation image (indicating faint or possibly unclear activity) that are not detected by the online algorithm (false negatives), or small partially visible processes detected by OnACID but ignored by the labelers ("false" positives).

**Figure 3:**
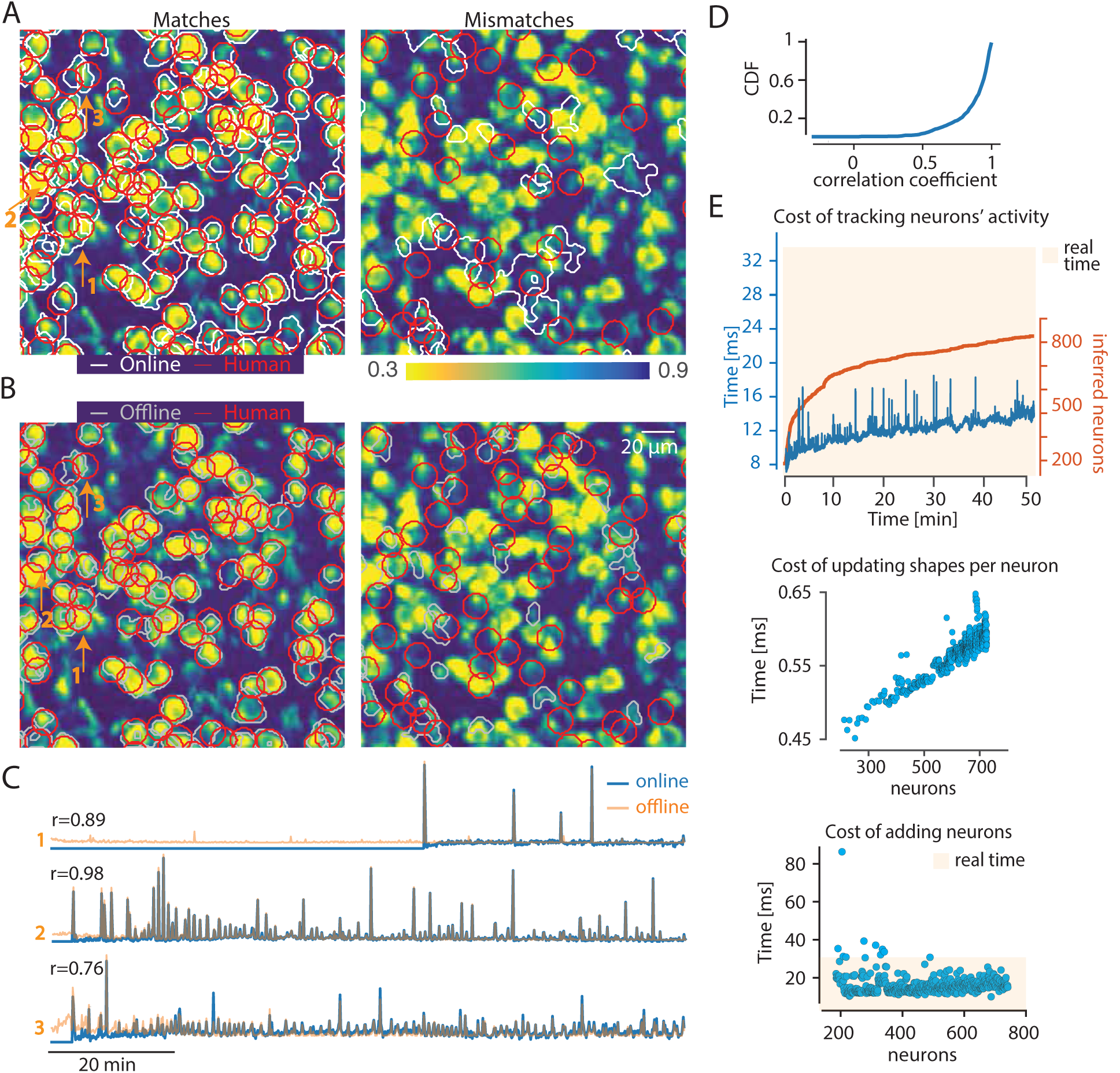
Application to an *in vivo* 50min long hippocampal dataset and comparison against an offline approach and manual annotation. A-left) Matched inferred locations between the online algorithm (white) and the manual annotation of Labeler 1 (red), superimposed on the max-correlation image. A-right) False positive (white) and false negative (red) mismatches between the online algorithm and a manual annotation. B) Same for the offline CNMF algorithm (grey) against the same manual annotation (red). The online approach outperforms the CNMF algorithm in the precision/recall framework (F_1_ score 0.77 vs 0.71). The images are restricted to a 200x200 pixel part of the FOV. Matches and non-matches for the whole FOV are shown in the supplement. C) Examples of inferred sources and their traces from the two algorithms and corresponding annotation for three indentified neurons (also shown with orange arrows in panels A,B left). The algorithm is capable of identifying new neurons once they become active, and track their activity similarly to offline approaches. D) Empirical CDF of correlation coefficients between the matched traces between the online and the offline approaches over the entire 50 minute traces. The majority of the correlation coefficients has very high values suggesting that the online algorithm accurately tracks the neural activity across time (see also correlation coefficients for the three examples shown in panel C). E) Timing of the online process. Top: Time required per frame when no shapes are updated and no neurons are updated (top). The algorithms is always faster than real time in tracking neurons and scales mildly with the number of neurons. Time required to update shapes per neuron (middle), and add new neurons (bottom) as a function of the number of neurons. Adding neurons is slower but occurs only sporadically affecting only mildly the required processing time (see text for details).

Fig. 3C shows examples of the traces from three selected neurons. OnACID can detect and track neurons with very sparse spiking over the course of the entire 50 minute experiment (Fig. 3C-top), and produce traces that are highly correlated with their offline counterparts. To examine the quality of the inferred traces (where ground truth collection at such scale is both very strenuous and severely impeded by the presence of background signals and neuropil activity), we compared the traces between the online algorithm and the CNMF approach on matched pairs of components. Fig. 3D shows the empirical cumulative distribution function (CDF) of the correlation coefficients from this comparison. The majority of the coefficients attain values close to 1, suggesting that the online algorithm can detect new neurons once they become active and then reliably track their activity.

**OnACID is faster than real time on average:** In addition to being more accurate, OnACID is also considerably faster as it required ∼27 minutes, i.e., ∼ 2× faster than real time on average, to analyze the full dataset (2 minutes for initialization and 25 for the online processing) as opposed to ∼1.5 hours for the offline approach and ∼10 hours for each of the annotators (who only select ROIs). Fig. 3E illustrates the time consumption of the various steps. In the majority of the frames where no spatial shapes are being updated and no new neurons are being incorporated, OnACID processing speed exceeds the data rate of 30Hz (Fig. 3E-top), and this processing time scales only mildly with the inclusion of new neurons. The cost of updating shapes and sufficient statistics per neuron is also very low (< 1ms), and only scales mildly with the number of existing neurons (Fig. 3E-middle). As argued before this cost can be distributed among all the frames while maintaining faster than real time processing rates. The expensive step appears when detecting and including one or possibly more new neurons in the algorithm (Fig. 3E-bottom). Although this occurs only sporadically, several speedups can be potentially employed here to achieve beyond real time at *every* frame (see also Discussion section), which would facilitate zero-lag closed-loop experiments.

**Application to *in vivo* 2p mouse parietal cortex data**: As a second application to 2p data we used a 116,000 frame dataset, taken at 30Hz over a 512×512 FOV (64min long). The first 3000 frames were used for initialization during which the CNMF algorithm found 442 neurons, before switching to OnACID, which by the end of the experiment found a total of 752 neurons (734 after filtering for size). Compared to two independent manual annotations of 928 and 875 ROIs respectively, OnACID achieved F_1_= 0.76,0.79 significantly outperforming CNMF (F_1_= 0.65,0.66 respectively). The matches and mismatches between OnACID and Labeler 1 on a 200x200 pixel part of the FOV are shown in Fig. 4A. Full FOV pairings as well as precision/recall metrics are given in Table 2.

**Figure 4:**
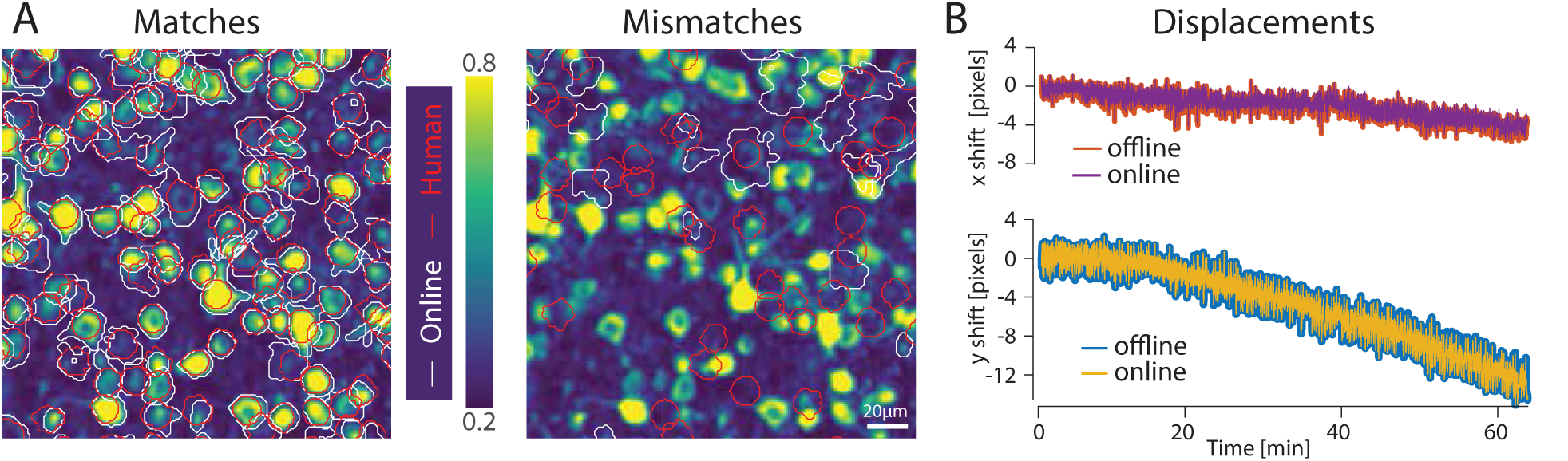
Application to an in *vivo* 64min long parietal cortex dataset. A-left) Matched inferred locations between the online algorithm (white) and the manual annotation of Labeler 1 (red). A-right) False positive (white) and false negative (red) mismatches between the online algorithm and a manual annotation. OnACID outperformed the offline CNMF algorithm. B) Displacement vectors estimated by OnACID during motion registration compared to a template based algorithm. OnACID estimates the same motion vectors at a sub-pixel resolution (see text for more details).

**Table 2:** Comparison of performance of OnACID and the CNMF algorithm using the precision/recall framework for the parietal cortex 116000 frames dataset. For each row-column pair, the column dataset is regarded as ground truth. The numbers in the parenthesis are the precision and recall, respectively, preceded by their harmonic mean (*F*_1_ score). OnACID significantly outperforms the offline CNMF approach when benchmarked against two independent manual annotations within the precision/recall framework that match highly with each other.

| $F_1$ (precision, recall) | Labeler 1 | Labeler 2 | CNMF |
| --- | --- | --- | --- |
| OnACID | 0.76 (0.86,0.68) | 0.79 (0.86,0.72) | 0.65 (0.55,0.82) |
| CNMF | 0.65 (0.70, 0.60) | 0.66 (0.74,0.59) | - |
| Labeler 2 | 0.89 (0.86,0.91) | - | - |

For this dataset, rigid motion correction was also performed according to the simple method of aligning each frame to the denoised (and registered) background from the previous frame. Fig. 4B shows that this approach produced strikingly similar results to an offline template based, rigid motion correction method [27]. The difference in the displacements produced by the two methods was less than 1 pixel for all 116,000 frames with standard deviations 0.11 and 0.12 pixel for the x and y directions, respectively. In terms of timing, OnACID processed the dataset in 48 minutes, again faster that real time on average. This also includes thetime needed for motion correction, which on average took 5ms per frame.

## 4 Discussion - Future Work

Our current implementation performs all processing serially. In principle, significant speed gains can be obtained by performing computations not needed at each timestep (updating shapes and sufficient statistics) or occur only sporadically (incorporating a new neuron) in a parallel thread with shared memory. Moreover, different online dictionary learning algorithms that do not require the solution of an inverse problem at each timestep can potentially further speed up our framework [18].

For detecting centroids of new sources OnACID examines a static image obtained by computing the variance across time of the spatially smoother residual buffer. While this approach works very well in practice it effectively favors shapes looking similar to a pre-defined Gaussian blob (when spatially smoothed). Different approaches for detecting neurons in static images can be possibly used here, e.g., [24], [2], [30].

Apart from facilitating closed-loop behavioral experiments and rapid general calcium imaging data analysis, our online pipeline can be potentially employed to future, optical-based, brain computer interfaces [6, 23] where high quality real-time processing is critical to their performance.

## Acknowledgments

We thank Sue Ann Koay, Jeff Gauthier and David Tank (Princeton University) for sharing their cortex and hippocampal data with us. We thank Lindsey Myers, Sonia Villani and Natalia Roumelioti for providing manual annotations. We thank Daniel Barabasi (Cold Spring Harbor Laboratory) for useful discussions. AG, DC, and EAP were internally funded by the Simons Foundation. Additional support was provided by SNSF P300P2_158428 (JF), and NIH BRAIN Initiative R01EB22913, DARPA N66001-15-C-4032, IARPA MICRONS D16PC00003 (LP).

## A Algorithmic description

Here we present in pseudocode the various steps of the online processing pipeline. For ease of exposition, some details and speedup tricks used in the actual implementation have been omitted.

Algorithms 2 and 3 describe the simple greedy procedure for partitioning the components into disjoint groups where the elements of each group do not overlap spatially with each other. This procedure is used for updating the traces of the neurons in vector form (Alg. 4) leading to substantial speed benefits. Algorithm 5 describes the procedure of detecting and screening possible new components. Finally, Algorithm 6 describes the process of updating the shapes, similar to [21].

### Algorithm 2 DetermineGroups

**Require:** Spatial components matrix *A*, number of components *K*

1: *G* =ø

2: **for** *i* = 1→ *K* **do**

3: *G* ←JoinGroups(A[:1 : *i* −1]; *G*; *i* − 1; a_*i*_)

4: **end for**

5:**return** *G*

### Algorithm 3 JoinGroups

**Require:** Spatial components matrix A, current groups G, number of components K, new component **a**

1:*N*_*G*_ = |*G*| ⊳number of groups

2:repeat = **True**

3:*g*←1

4:**while** repeat **do**

5:**if** ≤ *g* = *N*_*G*_ then

6 **if** a^┬^ a_l_= 0, ∀l ∊ *G*_*j*_ then ⊳Test for overlap with current group

7 *G*_*g*_ ←*G*_*g*_ ∪{*K* + 1}

8 repeat = **False**

9 else

10 *g* ← *g* + 1

11 **end if**

12 **else**

13 *N*_*G*_←*N*_*G*_+ 1

14 *G*_*NG*_ = {*K* + 1}⊳Create a new group

15 *G*←{*G*, *G*_*NG*_} ⊳ Add to list of groups

16 repeat = **False**

17 **end if**

18 **end while**

19 **return *G***

### Algorithm 4 UpdateTraces

**Require:** Spatial footprints matrix Ã = [A, b], current value of temporal traces ċ = [c; f], current data frame y, groups G, tolerance level ε

1 u = Ã ^┬^y

2 V = Ã TÃ

3 V = diag{V}

4 ċ _old_← 0

5 while ||ċ − ċ _old_||≥|| ċ _old_|| do

6 ċ _old_← ċ

7 for i = 1 → |G| do

8 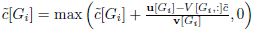 ⊳ (Division is pointwise)

9 **end for**

10 **end while**

11 **return** ċ

### Algorithm 5 DetectnewComponents

**Require:** Spatial footprints matrix [A, b], temporal traces matrix [C; f], current number of components K,current state of groups G, current residual buffer R_buf_, current data frame y, sufficient statistics W, M.Parameters: radius of Gaussian kernel T, threshold for correlation in space r_s_, threshold for correlation in time rt.

1 repeat = **True**

2 R_buf_← [R_buf_[:, 1 : l_b_− 1], y − [A, b][C; f][:, end]] ⊳ Update residual buffer

3 M_d_= MEDIAN (R_buf_)

4 ℝ_buf_← ℝ_buf_M_*d*_⊳ Subtract median along time for every pixel

5 *V* ← FILTER(ℝ_buf_, GAUSSIANKERNEL(τ)) ⊳ Filter residual in space

6 *E* ← ∑_*i*_*V*[:,*i*].^2^ ⊳Compute energy value for each pixel

7 **while** repeat **do**

8 (*i*_*x*_, *i*_*y*_) = arg max E ⊳ Find the point of maximum variance

9 *N*(*i*_*x*_,*i*_*y*_”) = {(*x*,*y*): |*x* − *i*_*x*_| ≤ τ, |y − *i*_*y*_| ≤ *t*} ⊳Define a neighborhood around (*i*_*x*_,*i*_*y*_)

10 [*a*_new_, *C*_new_] = NNMF(R_buf_[*N*(*i*_*x*_,*i*_*y*_),:], 1) ⊳ Perform a local rank-1 NMF

11 *r* = CORR(*a*_new_, MEAN(ℝ_buf_)) ⊳ Compute correlation coefficient in space

12 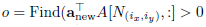 ⊳ Find components that overlap

13 if ∃*j*∊ o : CORR(cnew, *C*[*j*, *t* − *l*_*b*_+ 1 : *t*]) ⊳ *r*_*t*_ then

14 *r* ←0 ⊳ Detect possible duplicates and stop procedure

15 end if

16 if *r* > *r*_*s*_ then ⊳ New component is accepted

17 Zero-pad a_new_ and c_new_ to match dimensionality

18 *K* ← *K* + 1

19 *G* ← JOINGROUPS(*A*, *G*, anew)

20 *A* ← [*A*, *a*_new_]

21 *C* ← [*C*; *c*_new_]

22 ℝ_buf_← ℝ_buf_*a*_new_*c*_new_

23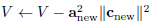

24 *W*, *M* ← UPDATESUFFSTATISTICS(*W*, *M*, *y*_*t*_, *c*_new_)⊳ Equation (5)

25 **Else**

26 repeat = **False**

27 **end if**

28 **end while**

29 **return** [A, b], [C, f], K, G, R**buf**, W, M

### Algorithm 6 UpdateShapes

**Require:** Sufficient statistics W, M, current value of spatial footprints Ã = [A, *b*], list of components to be updated l, maximum number of iterations iterations m_iter_

1 iter ← 0

2 **while** iter < m_iter_ **do**

3 **for** i ∊ l **do**

4 **p** = find(Ã [:, i] > 0) ⊳ Find the pixels where component i can be non-zero

5 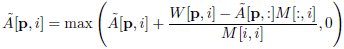

6 **end for**

7 iter ←iter + 1

8 **end while**

9 **return** Ã

## B Dataset Details

**Parietal cortex dataset**: Data was obtained from the parietal cortex of a transgenic GCaMP6f-expressing mouse during a behavioral task. Field of view was approximately 500x500 µm^2^ (512x512 pixels) in size and at depth 125µm below the dura surface. Horizontal scans of the laser were performed using a resonant galvanometer, resulting in a frame acquisition rate of 30Hz. More details can be found in [20].

**Hippocampal dataset**: Data was obtained from the hippocampus of a transgenic GP2.11 (Thyl- GCaMP3) mouse generated by the Janelia Farms GENIE Project (Jackson Labs, C57BL/6J-Tg(Thy1- GCaMP3)GP2.11Dkim/J). FOV was approximately 500x500 µm^2^, of size 512 x 512 pixels, cropped to 483 x 492 pixels after rigid registration and removal of empty border lines. Horizontal scans of the laser were performed using a resonant galvanometer, resulting in a frame acquisition rate of 30Hz. More details can be found in [13].

## C Supplementary Movie

**Evolution of the OnACID algorithm on toy simulated data:** Top. Raw movie (left). Denoised movie reconstructed from all components (middle). Noiseless ground truth (right). Bottom. Residual movie (left). Inferred (middle) and ground truth (right) spatial components. A 64x64 pixel FOV containing 35 artifical neurons was simulated for this example. Movie is truncated in time for space reasons.

## D Simulation details

We generated a dataset of size 256x256 pixels and duration T = 2000 frames containing N = 400 neurons. The neural centers 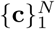were generated using a Halton sequence to cover the space uniform pseudo-randomly.

The unnormalized neural shapes were modeled as the difference of two 2D-Gaussians.

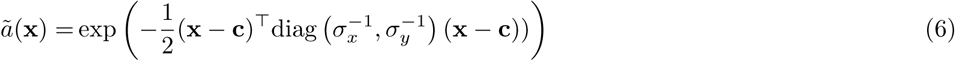

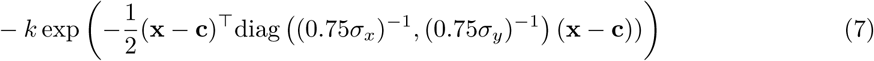

where x denotes the position of the considered pixel. To incorporate heterogeneity the standard-deviation *σ*_*x*_ and *σ*_*y*_ in x- and y-direction of the wider Gaussian was drawn i.i.d. uniform randomly from the interval [2.5, 3.5]. These values were multiplied by 0.75 to obtain the standard-deviation of the smaller subtracted Gaussian. The magnitude *k* of the subtracted Gaussian was drawn i.i.d uniform randomly from the interval [0.2,0.8] for each neuron.

The spike train s of each neuron was drawn from a homogeneous Poisson process. The neural firing rate was 0.5 Hz and the frame rate 30 Hz. The calcium traces C were obtained by convolving the spike trains S with an exponentially decaying kernel with time constant 1 s.

The background B was modeled as rank 1 term, where the temporal and spatial component were each drawn from a Kronecker Gaussian process with RBF kernel. The temporal length scale was 300 frames and the spatial length scale 50 pixels. Finally, the simulated raw data is the sum of background B and neural contribution AC corrupted by Gaussian noise, Y ? N(B + AC, 0.2^2^).

**Figure 5:**
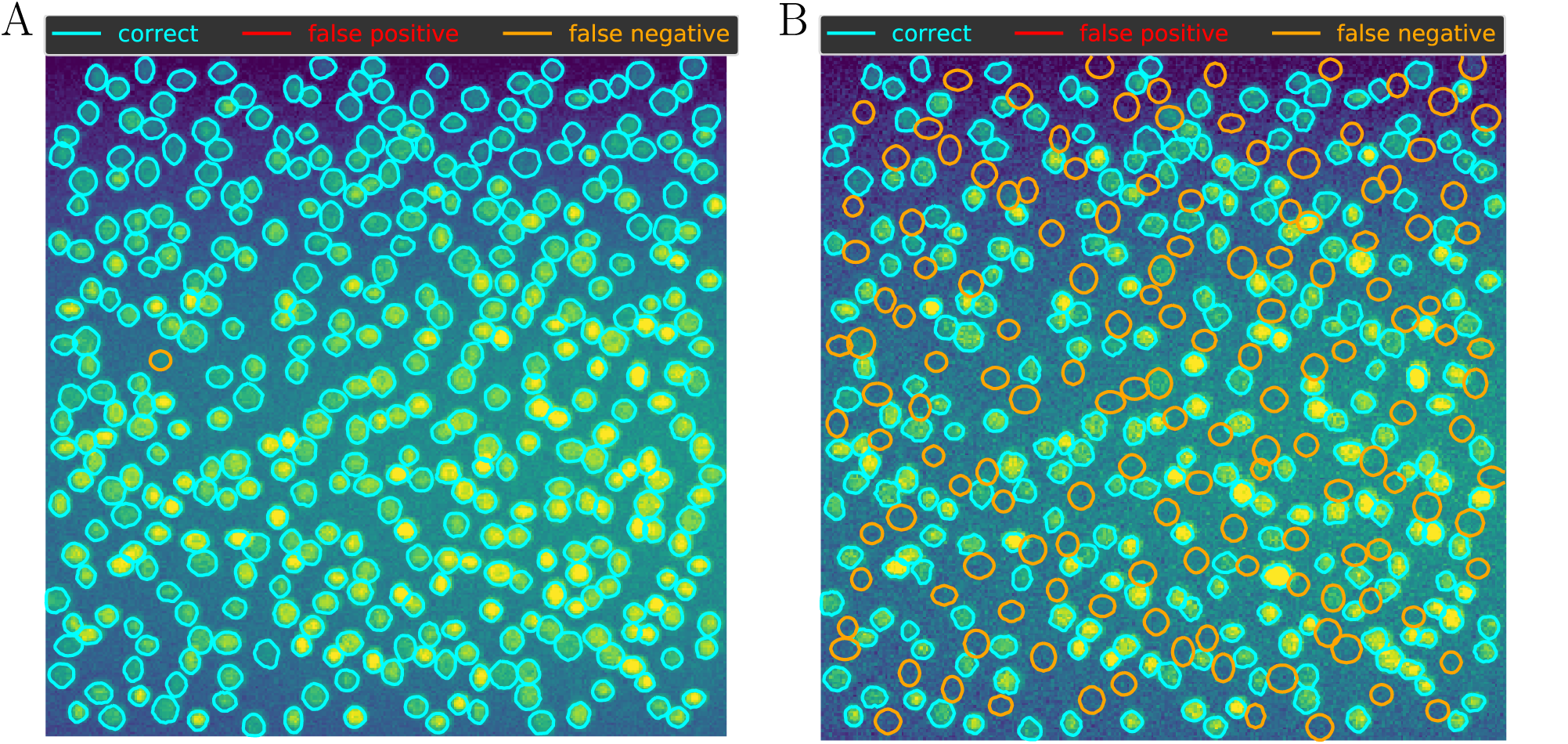
Simulated data. Detected components in batch mode. A) Full data B) Short initial batch.

**Figure 6:**
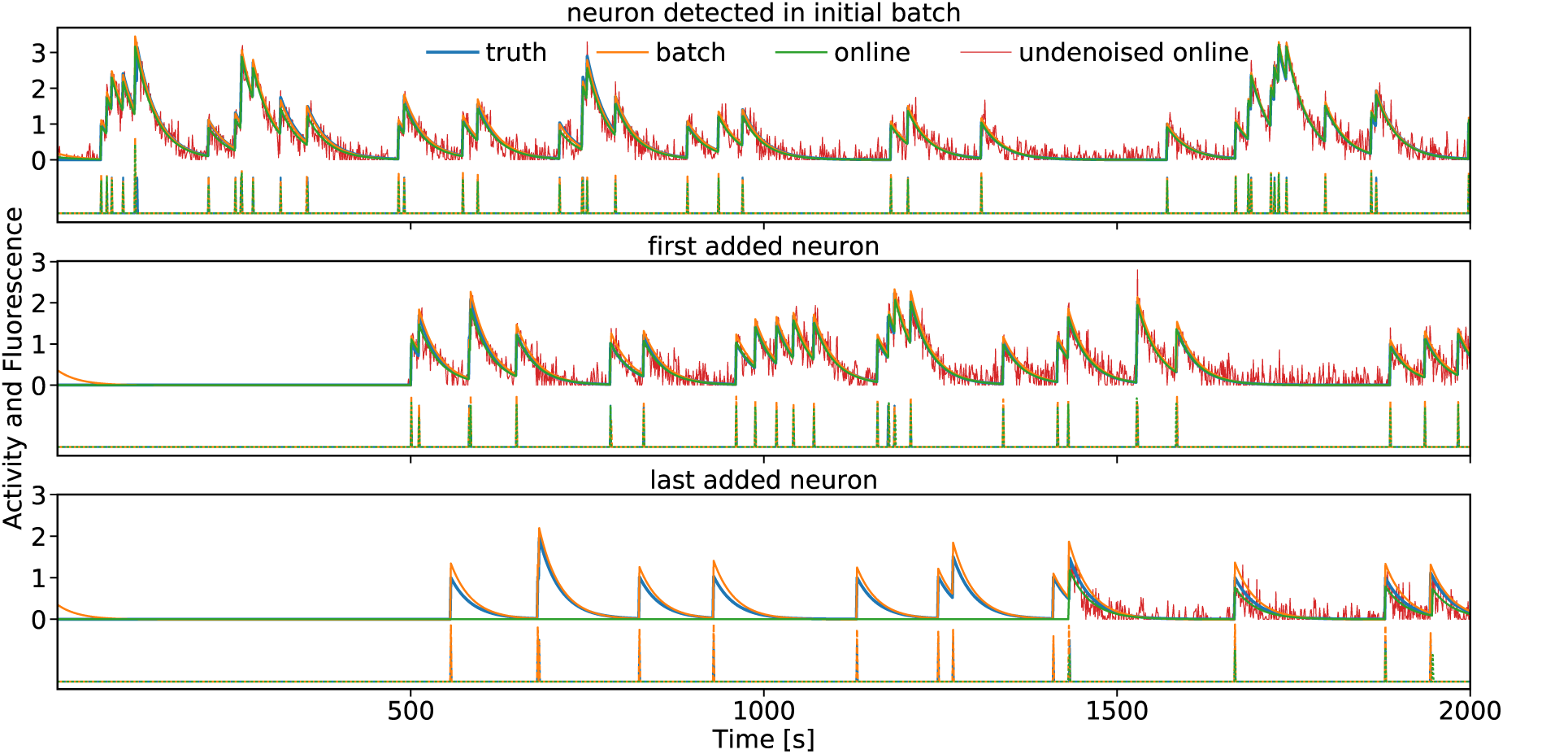
Simulated data. Traces of three neurons selected by time of detection. Upper traces show demixed (red), denoised (orange, green) and ground truth (blue) calcium fluorescence. Lower traces show deconvolved neural activity using the same coloring scheme.

## E Detailed comparison between OnACID and manual annotations for the hippocampal 2-photon dataset

In the following pages, Figures 7-12 show the detailed matches and mismatches between On ACID and the two manual annotations, as well as the two manual annotations against each other.

**Figure 7:**
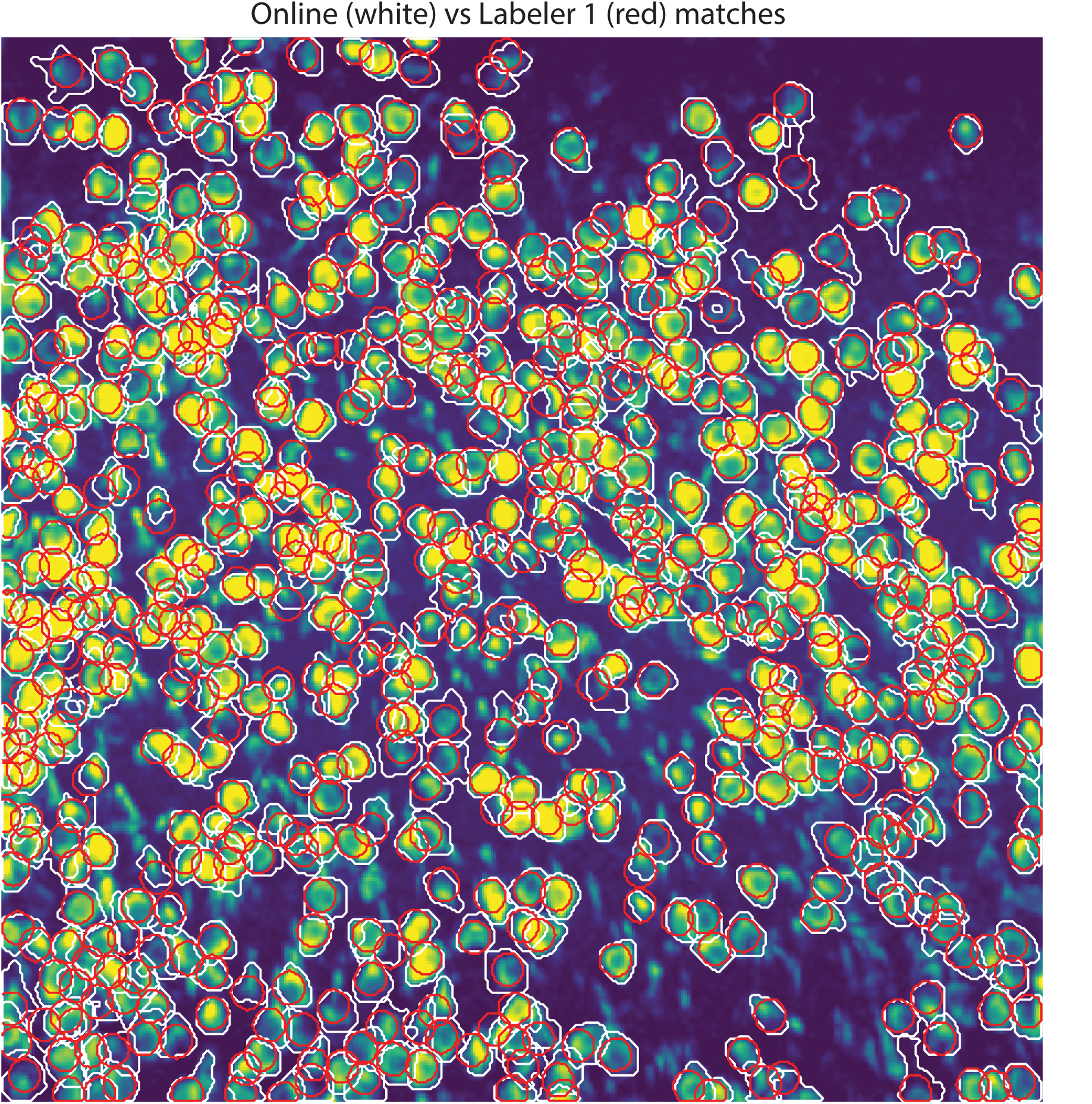
Matches between OnACID (white) and Labeler 1 (red).

**Figure 8:**
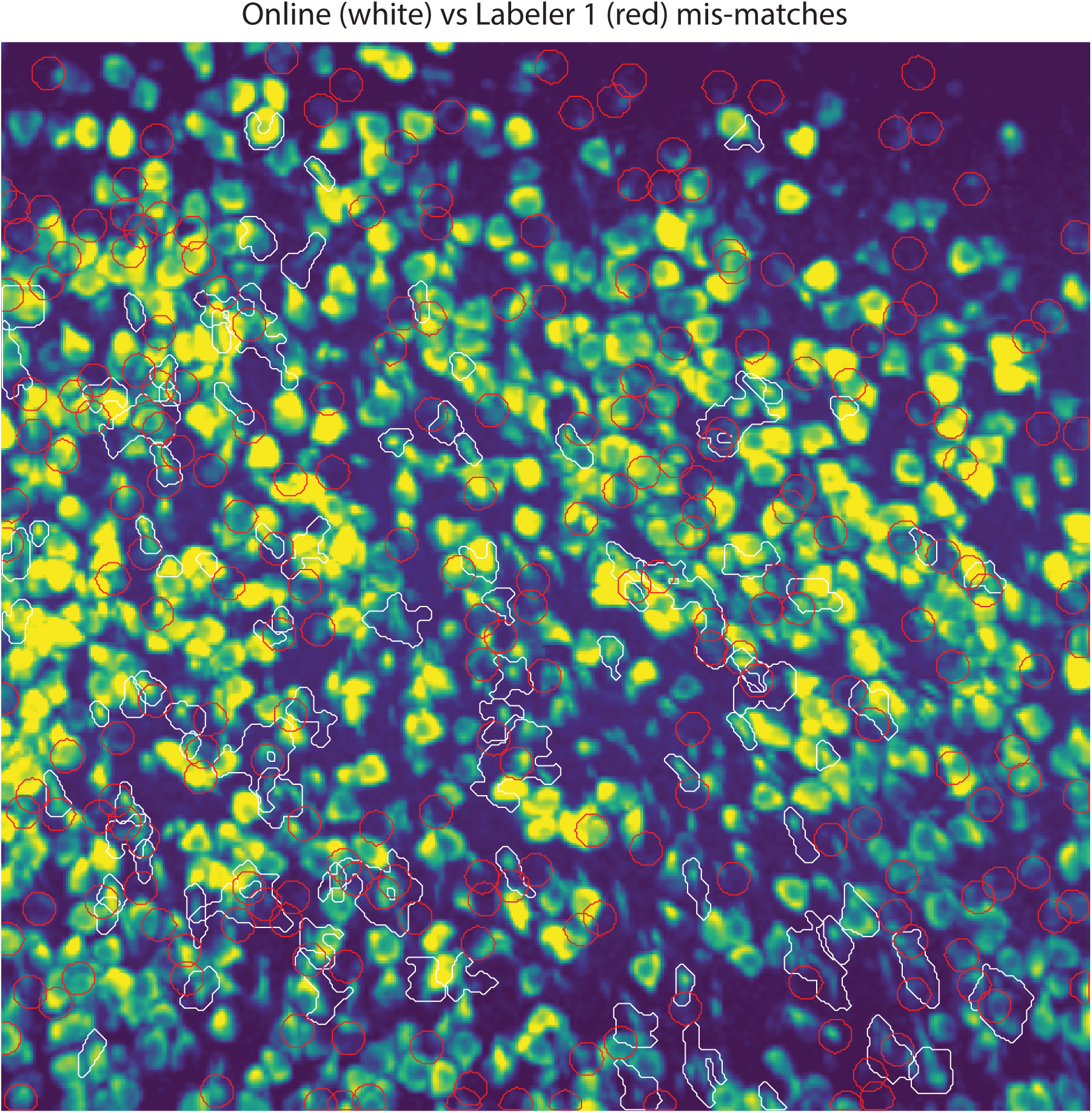
Mismatches between OnACID (white) and Labeler 1 (red).

**Figure 9:**
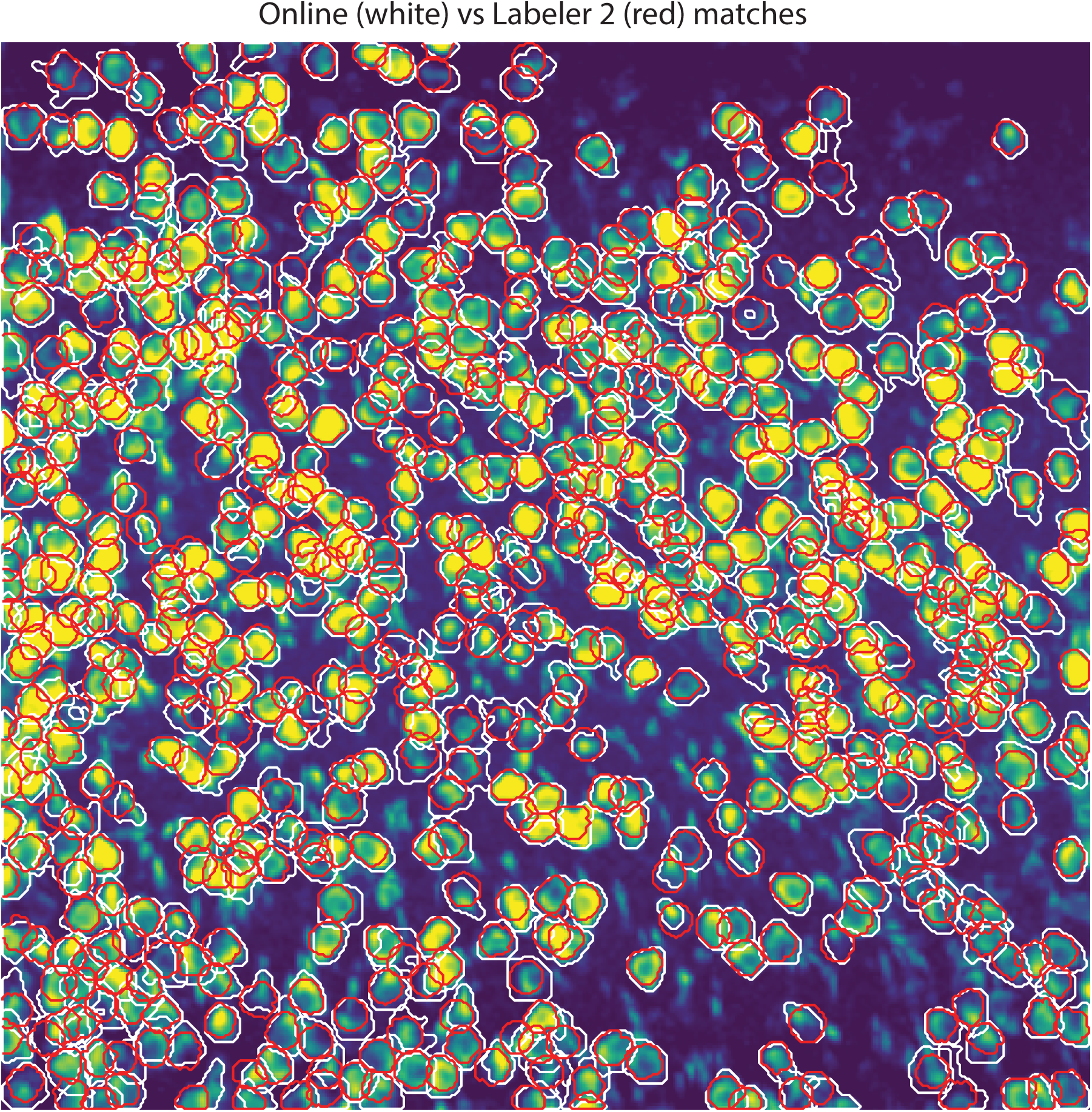
Matches between OnACID (white) and Labeler 2 (red).

**Figure 10:**
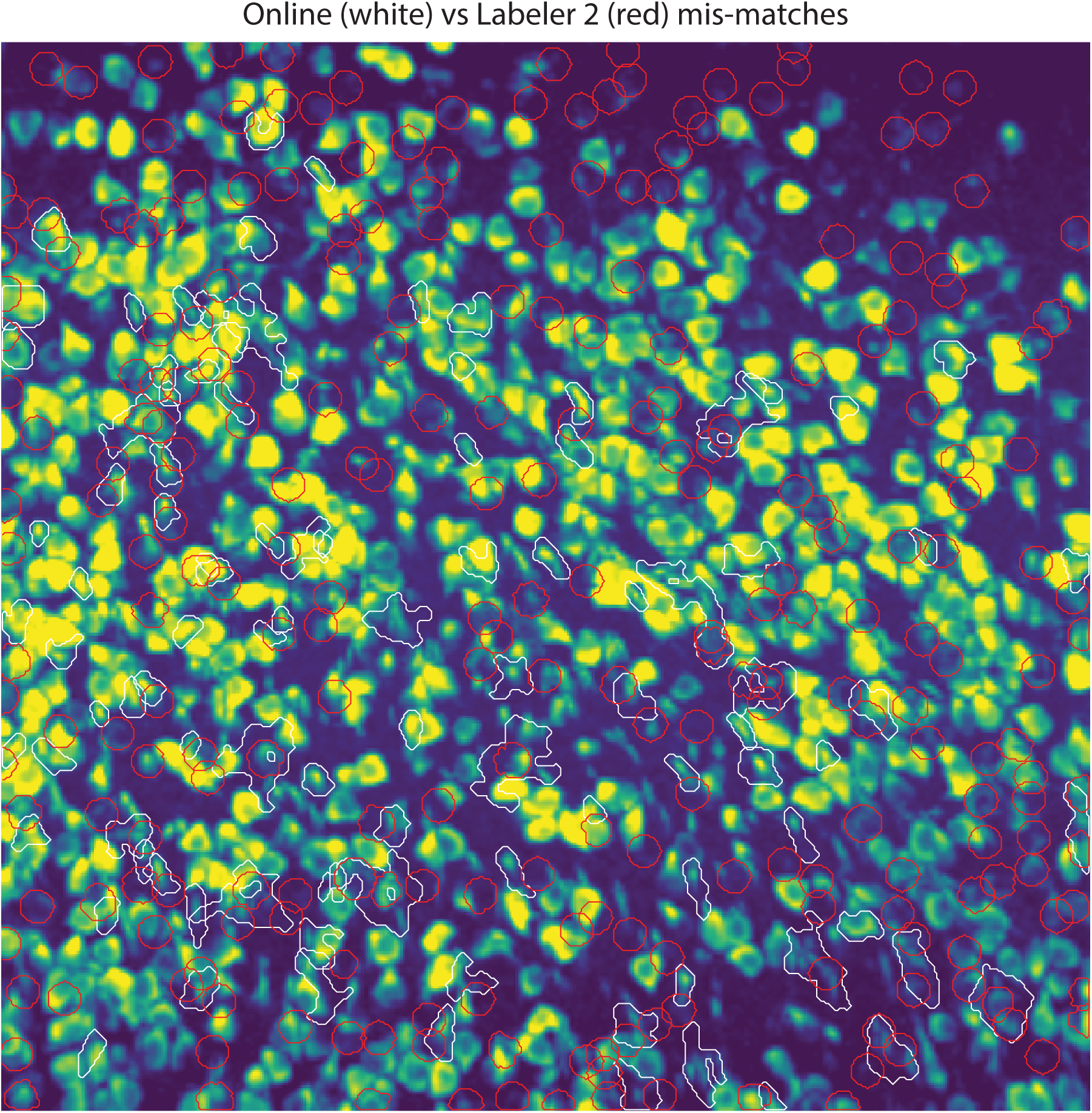
Mismatches between OnACID (white) and Labeler 2 (red).

**Figure 11:**
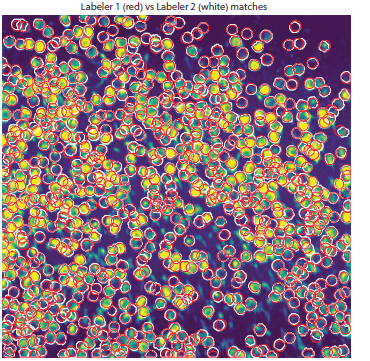
Matches between Labeler 1 (red) and Labeler 2 (white). The two labelers annotated the dataset independently and have a high degree of matching (F ^1^ = 0.89). The contour shapes are also similar for both annotators, as expected from the labeling process using the ImageJ Cell Wand tool.

**Figure 12:**
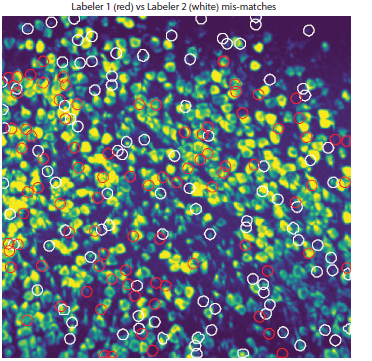
Mismatches between Labeler 1 (red) and Labeler 2 (white).

## F Detailed comparison between OnACID and manual annotations for the parietal cortex 2-photon dataset

In the following pages, Figures 13-18 show the detailed matches and mismatches between OnACID and the two manual annotations, as well as the two manual annotations against each other.

**Figure 13:**
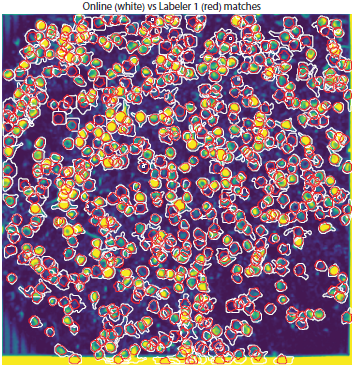
Matches between OnACID (white) and Labeler 1 (red).

**Figure 14:**
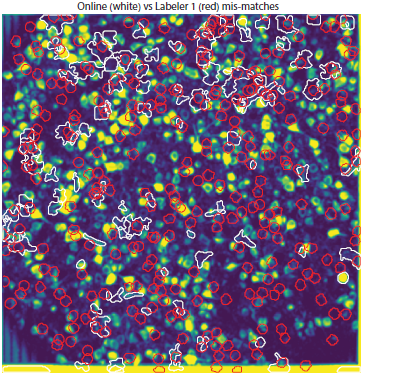
Mismatches between OnACID (white) and Labeler 1 (red).

**Figure 15:**
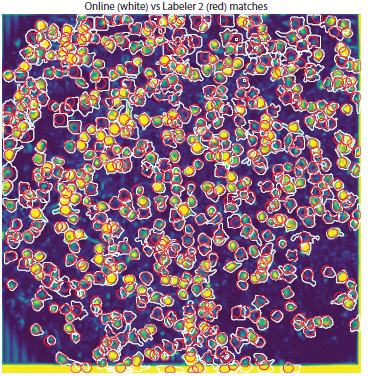
Matches between OnACID (white) and Labeler 2 (red).

**Figure 16:**
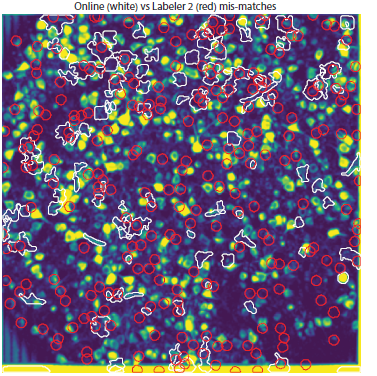
Mismatches between OnACID (white) and Labeler 2 (red)

**Figure 17:**
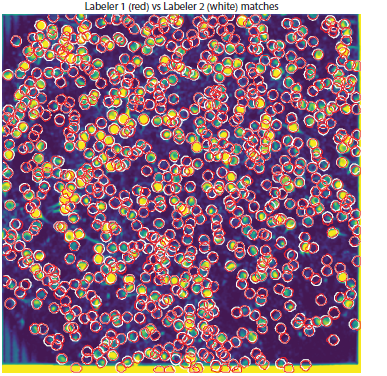
Matches between Labeler 1 (red) and Labeler 2 (white).

**Figure 18:**
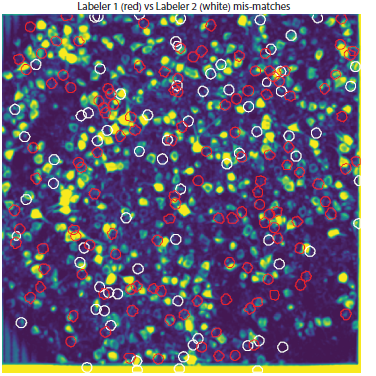
Mismatches between Labeler 1 (red) and Labeler 2 (white).

## Footnotes

1 The correlation image (CI) at every pixel is equal to the average temporal correlation coefficient between that pixel and its neighbors [29] (8 neighbors were used for our analysis). The max-correlation image is obtained by computing the CI for each batch of 1000 frames, and then taking the maximum over all these images.

2 Note that the Cell Magic Wand Tool by construction, tends to select circular ROI shapes whereas the results of the online algorithm do not pose restrictions on the shapes. As a result the computed Jaccard distances tend to be overestimated. This explains our choice of a seemingly high mismatch threshold.

## References

[1] Misha B Ahrens, Michael B Orger, Drew N Robson, Jennifer M Li, and Philipp J Keller. Whole-brain functional imaging at cellular resolution using light-sheet microscopy. Nature methods, 10(5):413–420, (2013).

[2] Noah Apthorpe, Alexander Riordan, Robert Aguilar, Jan Homann, Yi Gu, David Tank, and H Sebastian Seung. Automatic neuron detection in calcium imaging data using convolutional networks. In Advances in Neural Information Processing Systems, pages 3270–3278, 2016.

[3] Richard E Barlow, David J Bartholomew, JM Bremner, and H Daniel Brunk. Statistical inference under order restrictions: The theory and application of isotonic regression. Wiley New York, 1972.

[4] Matthew B Bouchard, Venkatakaushik Voleti, César S Mendes, Clay Lacefield, Wesley B Grueber, Richard S Mann, Randy M Bruno, and Elizabeth MC Hillman. Swept confocally-aligned planar excitation (scape) microscopy for high-speed volumetric imaging of behaving organisms. Nature photonics, 9(2):113–119, 2015.

[5] Edward S Boyden, Feng Zhang, Ernst Bamberg, Georg Nagel, and Karl Deisseroth. Millisecond-timescale, genetically targeted optical control of neural activity. Nature neuroscience, 8(9):1263–1268, (2005).

[6] Kelly B Clancy, Aaron C Koralek, Rui M Costa, Daniel E Feldman, and Jose M Carmena. Volitional modulation of optically recorded calcium signals during neuroprosthetic learning. Nature neuroscience, 17(6):807–809, (2014).

[7] Daniel A Dombeck, Anton N Khabbaz, Forrest Collman, Thomas L Adelman, and David W Tank. Imaging large-scale neural activity with cellular resolution in awake, mobile mice. Neuron, 56(1):43–57, (2007).

[8] Valentina Emiliani, Adam E Cohen, Karl Deisseroth, and Michael Hausser. All-optical interrogation of neural circuits. Journal of Neuroscience, 35(41):13917–13926, (2015).

[9] Jeremy Freeman, Nikita Vladimirov, Takashi Kawashima, Yu Mu, Nicholas J Sofroniew, Davis V Bennett, Joshua Rosen, Chao-Tsung Yang, Loren L Looger, and Misha B Ahrens.Mapping brain activity at scale with cluster computing. Nature methods, 11(9):941–950, (2014).

[10] Johannes Friedrich, Weijian Yang, Daniel Soudry, Yu Mu, Misha B Ahrens, Rafael Yuste, Darcy S Peterka, and Liam Paninski. Multi-scale approaches for high-speed imaging and analysis of large neural populations. bioRxiv, page 091132, 2016.

[11] Johannes Friedrich, Pengcheng Zhou, and Liam Paninski. Fast online deconvolution of calcium imaging data. PLOS Computational Biology, 13(3):e1005423, 2017.

[12] Sahil Garg, Irina Rish, Guillermo Cecchi, and Aurelie Lozano. Neurogenesis-inspired dictionary learning: Online model adaption in a changing world. arXiv preprint arXiv:1701.06106, 2017.

[13] J. Gauthier and D.W. Tank. A subset of ca1 and subiculum neurons selectively encode rewarded locations. In Computational and Systems Neuroscience Meeting, Cosyne, 2016.

[14] A Giovannucci, J Friedrich, B Deverett, V Staneva, D Chklovskii, and E Pnevmatikakis. Caiman: An open source toolbox for large scale calcium imaging data analysis on standalone machines. Cosyne Abstracts, 2017.

[15] David S Greenberg and Jason ND Kerr. Automated correction of fast motion artifacts for two-photon imaging of awake animals. Journal of neuroscience methods, 176(1):1–15, (2009).

[16] Logan Grosenick, James H Marshel, and Karl Deisseroth. Closed-loop and activity-guided optogenetic control. Neuron, 86(1):106–139, (2015).

[17] Manuel Guizar-Sicairos, Samuel T Thurman, and James R Fienup. Efficient subpixel image registration algorithms. Optics letters, 33(2):156–158, (2008).

[18] Tao Hu, Cengiz Pehlevan, and Dmitri B Chklovskii. A hebbian/anti-hebbian network for online sparse dictionary learning derived from symmetric matrix factorization. In Signals, Systems and Computers, 2014 48th Asilomar Conference on, pages 613–619. IEEE, 2014.

[19] Patrick Kaifosh, Jeffrey D Zaremba, Nathan B Danielson, and Attila Losonczy. Sima: Python software for analysis of dynamic fluorescence imaging data. Frontiers in neuroinformatics, 8:80, 2014.

[20] Sue Ann Koay, Ben Engelhard, Lucas Pinto, Ben Deverett, Stephan Thiberge, Carlos Brody, and David Tank. Neural dynamics in a mouse navigation and accumulation of visual evidence task. In Society for Neuroscience, number 739.07, 2016.

[21] Julien Mairal, Francis Bach, Jean Ponce, and Guillermo Sapiro. Online learning for matrix factorization and sparse coding. Journal of Machine Learning Research, 11(Jan):19–60, (2010).

[22] Eran A Mukamel, Axel Nimmerjahn, and Mark J Schnitzer. Automated analysis of cellular signals from large-scale calcium imaging data. Neuron, 63(6):747–760, (2009).

[23] Daniel J O'shea, Eric Trautmann, Chandramouli Chandrasekaran, Sergey Stavisky, Jonathan C Kao, Maneesh Sahani, Stephen Ryu, Karl Deisseroth, and Krishna V Shenoy.The need for calcium imaging in nonhuman primates: New motor neuroscience and brain-machine interfaces. Experimental neurology, 287:437–451, (2017).

[24] Marius Pachitariu, Adam M Packer, Noah Pettit, Henry Dalgleish, Michael Hausser, and Maneesh Sahani. Extracting regions of interest from biological images with convolutional sparse block coding. In Advances in Neural Information Processing Systems, pages 1745–1753, 2013.

[25] Marius Pachitariu, Carsen Stringer, Sylvia Schroder, Mario Dipoppa, L Federico Rossi, Matteo Carandini, and Kenneth D Harris. Suite2p: beyond 10,000 neurons with standard two-photon microscopy. bioRxiv, page 061507, 2016.

[26] Adam M Packer, Lloyd E Russell, Henry WP Dalgleish, and Michael Hausser. Simultaneous all-optical manipulation and recording of neural circuit activity with cellular resolution in vivo. Nature Methods, 12(2):140–146, (2015).

[27] Eftychios A Pnevmatikakis and Andrea Giovannucci. Normcorre: An online algorithm for piecewise rigid motion correction of calcium imaging data. bioRxiv, page 108514, 2017.

[28] Eftychios A Pnevmatikakis, Daniel Soudry, Yuanjun Gao, Timothy A Machado, Josh Merel, David Pfau, Thomas Reardon, Yu Mu, Clay Lacefield, Weijian Yang, et al. Simultaneous denoising, deconvolution, and demixing of calcium imaging data. Neuron, 89(2):285–299, (2016).

[29] Spencer L Smith and Michael Hausser. Parallel processing of visual space by neighboring neurons in mouse visual cortex. Nature neuroscience, 13(9):1144–1149, (2010).

[30] Quico Spaen, Dorit S Hochbaum, and Roberto Asin-Acha. HNCcorr: A novel combinatorial approach for cell identification in calcium-imaging movies. arXiv preprint arXiv:1703.01999, 2017.

[31] Joshua T Vogelstein, Adam M Packer, Timothy A Machado, Tanya Sippy, Baktash Babadi, Rafael Yuste, and Liam Paninski. Fast nonnegative deconvolution for spike train inference from population calcium imaging. Journal of neurophysiology, 104(6):3691–3704, (2010).

[32] Theo Walker. Cell magic wand tool, 2014.

